# Heterogeneous expression of Pil3 pilus is critical for *Streptococcus gallolyticus* translocation across polarized colonic epithelial monolayers

**DOI:** 10.1101/682278

**Authors:** Mariana Martins, Laurence du Merle, Patrick Trieu-Cuot, Shaynoor Dramsi

**Affiliations:** Department of Microbiology, Biology of Gram-positive Pathogens Unit, Institut Pasteur, Paris, France; Centre National de la Recherche Scientifique (CNRS) ERL 6002 Paris, France

**Keywords:** Pil3 pilus, *S. gallolyticus*, colon, translocation

## Abstract

*Streptococcus gallolyticus subspecies gallolyticus* (*Sgg*) is an opportunistic pathogen responsible for septicaemia and endocarditis in elderly persons. *Sgg* is also a commensal of the human gastrointestinal tract. Here we demonstrate that *Sgg* strain UCN34 translocates across tight intestinal barriers *in vitro* in a Pil3-dependent manner. Confocal images of UCN34 passage across human colonic cells reveals that *Sgg* utilizes a paracellular pathway. Pil3 was previously shown to be expressed heterogeneously and WT UCN34 consists of about 90% of Pil3_low_ and 10% of Pil3_high_ cells. We found that both the Δ*pil3* mutant and the Pil3+ overexpressing variant could not translocate across Caco-2 and T84 barriers. Interestingly, combining live Δ*pil3* mutant cells with fixed Pil3+ variants in a 10:1 ratio (mimicking UCN34 WT population) allowed efficient translocation of the Δ*pil3* mutant. These experiments demonstrate that heterogeneous expression of Pil3 plays a key role in optimal translocation of *Sgg* across the intestinal barrier.

**ABSTRACT IMPORTANCE:** *Streptococcus gallolyticus subsp. gallolyticus* (*Sgg*) is an opportunistic pathogen responsible for septicemia and infective endocarditis in elderly persons. *Sgg* is a commensal of the rumen of herbivores and transmission to humans most probably occurs through the oral route. In this work, we have studied how this bacterium crosses the intestinal barrier using well-known *in vitro* models. Confocal microscopy images revealed that *Sgg* UCN34 can traverse the epithelial monolayer in between adjacent cells. We next showed that passage of *Sgg* from the apical to the basolateral compartment is dependent on the heterogenous expression of the Pil3 pilus at the bacterial surface. We hypothesize that Pil3_high_ cocci adhere firmly to epithelial cells to activate transient opening of tight junctions thereby allowing the traversal of Pil3l*ow* bacteria.

## INTRODUCTION

*Streptococcus gallolyticus subsp. gallolyticus (Sgg)*, formerly known as *S. bovis* biotype I, is an emerging opportunistic pathogen responsible for septicemia and infective endocarditis in elderly and immunocompromised individuals (1, 2). This Gram-positive coccus is one of the few intestinal bacteria that have been consistently linked to colorectal cancer (CRC) over the last 40 years (3) and a recent study confirms this suspicion (4). Whether *Sgg* is a driver and/or passenger of CRC is the subject of recent studies and evidence for both models have been obtained experimentally (5–10).

The first complete *S. gallolyticus* genome obtained was that of strain UCN34, isolated from a patient suffering from endocarditis and later diagnosed for colon cancer, provided important insights on the adaptation and virulence strategies developed by this bacterium (11). We previously showed that the Pil1 pilus is important for binding to collagen through the pilus associated adhesin Pil1A as well as colonization of heart valves in a rat model of experimental endocarditis (12). We also showed that the Pil3 pilus is important for colonization of the host colon via attachment to the intestinal mucus covering colonic cells (13, 14).

Pil1 and Pil3 are both expressed heterogeneously in the UCN34 population, with a majority of cells weakly piliated and a minority highly piliated, through a mechanism combining phase variation and attenuation (15).

In this report we investigated how *Sgg* adheres to and translocates across tight epithelial barriers in the absence of a secreted mucus layer. We demonstrate a key role of the Pil3 pilus in this process and visualized *Sgg* passage through a paracellular pathway. Our results indicate that in the UCN34 WT *Sgg* population, the highly piliated bacteria activate opening of the tight junctions to allow paracellular crossing of Pil3_low_ expressing bacteria. This demonstrates the functional relevance of Pil3 heterogeneity by a very unique mechanism.

## RESULTS

### The Sgg Pil3 pilus enhances adherence to human colonic cells

Streptococcal pili have been implicated both in adherence to eukaryotic cells as well as in bacterial translocation across host epithelial barriers (16, 17). We previously showed that the Sgg Pil3 pilus strongly contributes to bacterial attachment to human mucus-producing cells HT29-MTX (13). Therefore, we wondered whether Pil3 could also play a role in bacterial translocation. However, HT29-MTX cells were not able to form tight epithelial barriers on Transwell filters. We therefore tested two other well-studied human colonic cell lines Caco-2 and T84, which are both able to form tightly polarized epithelial monolayers *in vitro* (18).

We first compared adherence of *Sgg* UCN34 (WT), a highly Pil3 piliated variant (Pil3+) and a deletion mutant (Δ*pil3*) to Caco-2 cells at different time points. As shown in Fig. 1, higher Pil3 expression levels led to increased bacterial adhesion to Caco-2 cells as compared to the WT while bacterial adherence in the absence of Pil3 was decreased at 6h post-infection. Very similar results were observed in T84 cells (**Fig. S2 A**). It is worth noting that *Sgg* strain UCN34, which displays intermediate adherence, is composed of a heterogeneous population with approximately 10-20% highly expressing Pil3 pilus (Pil3^high^) and 80-90% weakly piliated (Pil3^low^). Taken together, these results indicate that the Pil3 pilus also contributes to *Sgg* adhesion to human colonic epithelial cells. We next investigated Sgg translocation in these model cell lines.

**Fig. 1.**
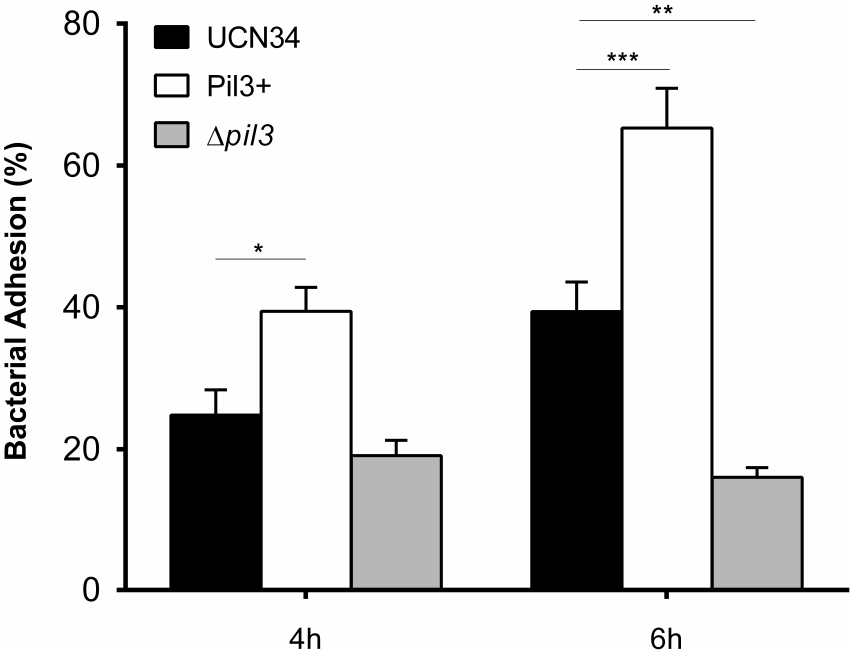
Role of Pil3 pilus in the adherence of *Sgg* UCN34 to human colorectal cancer cells Caco-2. Adherence is presented as percentage of bacterial inoculum after 4h and 6h at 37°C at a multiplicity of infection of 10 bacteria per cell. Planktonic growth of the various *Sgg* strains in this cell medium was monitored and did not change during the infection period.

### *S. gallolyticus* translocates across colonic epithelial barriers

In order to study translocation of <u>*Sgg*</u>, we first established an *in vitro* model of polarized cells using human colonic Caco-2 and T84 cell lines (**Fig. S1 and S2**). In particular, the T84 cell line has been widely used because of its ability to form a tall columnar epithelial monolayer and to develop tight intercellular junctions when grown on a permeable support, with the basolateral surface attached to the support and the apical surface exposed. Both cell lines were cultured on Transwell inserts for 3 to 21 days which allowed their polarization and differentiation. On these filters, fully differentiated Caco-2 and T84 cells expressed well-organized cell-to-cell junctions, forming a cell monolayer mimicking the intestinal epithelial barrier. Integrity of the epithelial barrier was monitored over time using two complementary methods: transepithelial electrical resistance (TER) of the monolayers and permeability to 4 kDa FITC-Dextran molecular ruler (**Fig. S1**). After 11 days in culture, the Caco-2 and T84 monolayers were already impermeable to the 4 kDa dextran molecules and reached a TER of approximately 300 and 2500 Ω cm^2^, respectively (**Fig. S1**). The capacity of *Sgg* UCN34 to translocate across the impermeable Caco-2 and T84 barriers was assessed at days 7, 14, and 21 and although translocation of UCN34 WT increased with cell differentiation (data not shown), day 14 was chosen as the time point giving the most consistent results. In order to demonstrate that bacterial translocation is an active process, cell monolayers were apically infected with *Sgg* UCN34 by inverting the transwell (lower chamber of the transwell insert) and translocated bacteria were recovered in the upper chamber at 2, 4, and 6 h post-infection (**Fig. 2A**). The rate of translocation increased over time with a maximum translocation of about 10% in Caco-2 (**Fig. 2B**) and 12% in T84 monolayers (**Fig. S2B**) at 6h post-infection. The effect of *Sgg* on epithelial barrier function was also assessed by measuring the TER following infection. TER values remained stable upon infection and were comparable to non-infected control monolayers, indicating no major disruption of the epithelial barrier by *Sgg* UCN34 (**Fig. S1C and Fig. S2D**). In order to visualize bacteria during the translocation process, confocal imaging was carried out on infected Caco-2 cells grown on filters at 6 h post-infection. Caco-2 cells were stained with E-cadherin which localizes at epithelial junctions, actin was stained with phalloidin to visualize the cytoskeleton and bacteria were labeled using a specific polyclonal antibody directed against strain UCN34. During translocation Sgg UCN34 was found primarily close to E-cadherin, at epithelial junctions between adjacent cells (**Fig. 2C**). These images suggest that UCN34 uses a paracellular route with a transient opening of cell junctions, as suggested by a previous study (3). Cell monolayers were considered well polarized, as demonstrated by the specific accumulation of actin close to the apical membrane of the cells. No intracellular *Sgg* UCN34 could be detected inside Caco-2 cells (data not shown). Thus, these results strongly suggest that *Sgg* UCN34 is able to translocate epithelial barriers of human intestinal cells through a process involving a transient and subtle opening of tight junctions.

**Fig. 2.**
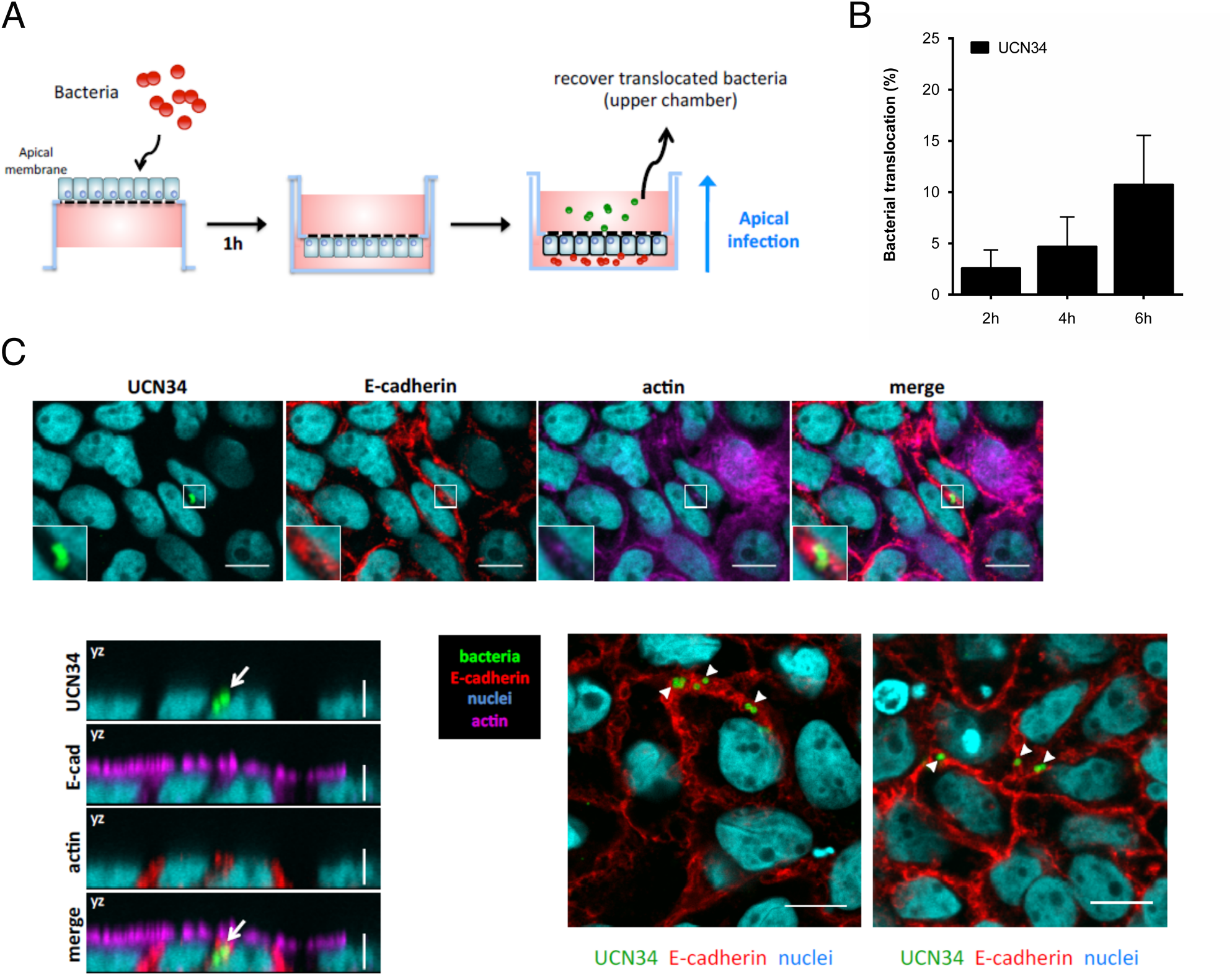
Translocation of *Sgg* UCN34 across Caco-2 cells. A- Schematic representation of the infection process using inverted transwell system. The transwell inserts were washed, inverted and incubated for about 1h to allow bacterial attachment to the cells. Then the inserts were placed back into a new 6 well plate. At each time point of infection, the medium for the upper compartment was completely recovered for CFU quantification and replaced by fresh medium. B- Translocation of *Sgg* UCN34 across Caco-2 monolayer. Cells were infected for 2, 4 and 6 h with a multiplicity of infection of 10 bacteria per cell. Translocation values are relative to the inoculum and represent 5 independent experiments performed in duplicate. C- Visualization of *Sgg* UCN34 translocation using confocal microscopy. After 6 h of infection, Caco-2 monolayer were fixed, permeabilized and stained with a monoclonal antibody against E-cadherin (in red), A547 conjugated-phalloidin to visualize actin (purple), and Hoescht 33342 to reveal cells nuclei (cyan). Bacteria were detected with a specific polyclonal antibody raised against the whole bacterium (green). The upper panels show an YX view of the filter, whereas the lower panels show an YZ view. The arrows are pointing to UCN34. In the upper panels the scale bar represents 10 μm and in the yz planes 5 μm. Right panel: representative image from another independent experiment. The scale bar corresponds to 10 μm.

### Pil3 pilus heterogeneity in *Sgg* is required for efficient bacterial translocation

To gain further mechanistic insights about *Sgg* translocation across epithelial barriers, we analyzed the possible contribution of the Pil3 pilus in this process. Caco-2 (**Fig. 3A**) and also T84 (**Fig. S2B**) monolayers of cells were infected with *Sgg* UCN34 (WT), a Pil3+ variant and the Δ*pil3* mutant for 2, 4, and 6h. WT *Sgg* was able to translocate these barriers at a rate of 5-10%. In contrast, the otherwise isogenic Pil3+ variant and Δ*pil3* mutant were significantly impaired by about 5-fold in this process as compared to WT UCN34. These results demonstrate that: i) the Pil3 pilus is essential for *Sgg* translocation across intestinal barriers as the Δ*pil3* mutant is unable to translocate and more surprisingly ii) that Pil3 pilus heterogeneity is also functionally important since a highly expressing Pil3+ variant cannot translocate these cell monolayers. We hypothesize that increased adherence of the Pil3+ variant at the apical surface of the cells impaired bacterial translocation. Hence, it appears that a delicate balance in Pil3 expression is important for efficient *Sgg* translocation. To test this hypothesis, we artificially mimicked the expression levels of the Pil3 pilus in the natural *Sgg* UCN34 population (around 90% of Pil3_low_ and 10% of Pil3_high_). To achieve this, we mixed the Δ*pil3* mutant with PFA killed Pil3+ variant in a proportion of 9:1, respectively. We then infected Caco-2 monolayers with this mixture and observed the capacity of the live Δ*pil3* mutant to translocate (**Fig. 3B**). Interestingly, the presence of fixed Pil3+ bacteria allowed the Δ*pil3* mutant to cross the Caco-2 monolayer with the same efficiency as *Sgg* UCN34. These data demonstrate that heterogeneous expression of Pil3 in *Sgg* is crucial for efficient translocation across intestinal barriers. Furthermore, blocking experiments were carried out in this experimental setting using specific antibodies against Pil3 (**Fig. 3C**). First, we showed that the addition of antibodies against both Pil3 subunits impaired translocation of the Δ*pil3* mutant in the presence of PFA killed Pil3+ variant cells (**Fig. 3C**). The Pil3 pilus is composed of a putative tip-located Pil3A adhesin and of a major Pil3B pilin subunit constituting the backbone of the Pil3 filamentous structure. In order to demonstrate the specific role of the Pil3A adhesin in *Sgg* translocation, antibodies against Pil3A were tested in the same experimental setting and were shown to be sufficient to block translocation of the Δ*pil3* mutant with PFA killed Pil3+ cells (**Fig. 3D**). As a control, we used a similar type of antibody raised against Pil1 that did not prevent translocation of live Δ*pil3* mutant cells in the presence of PFA killed Pil3+ bacteria. Altogether, these results demonstrate that heterogeneous expression of the Pil3 pilus in the *Sgg* UCN34 is critical for its ability to translocate efficiently across tight epithelial barriers, and that the paracellular opening process depends on Pil3A adhesin interaction with one or several as yet unknown host cell receptor(s).

**Fig 3.**
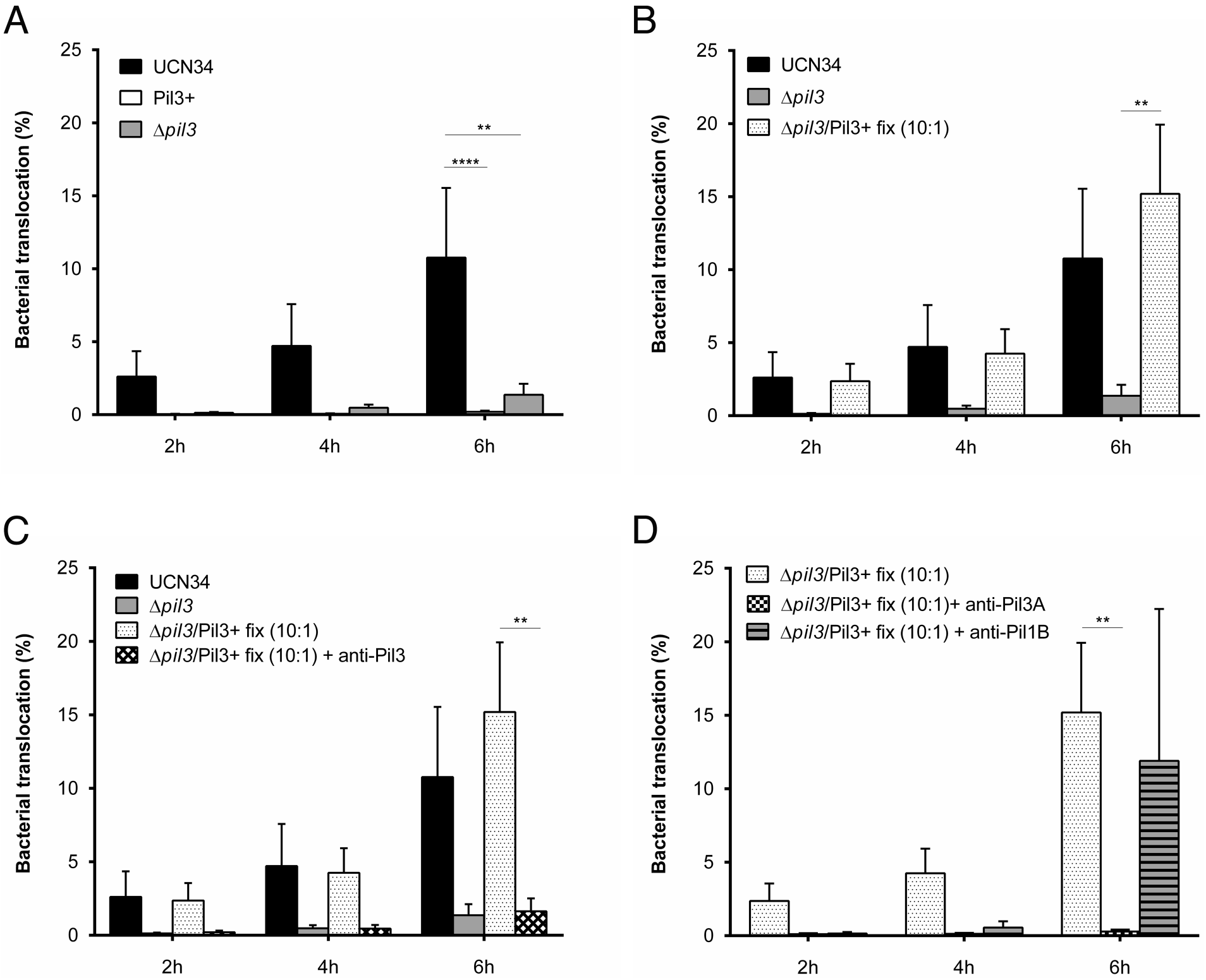
Bacterial translocation across Caco-2 monolayer is dependent on Pil3 pilus heterogeneous expression. A- UCN34, Pil3+ and Δ*pil3* translocation across Caco-2 monolayer after 2 h, 4 h and 6 h of infection. B- translocation of a mix of 10:1 Δ*pil3* and fixed killed Pil3+ as compared to WT UCN34 and Δ*pil3* alone. C- inhibition of translocation with antibodies against Pil3. D- inhibition of translocation with specific antibodies directed against Pil3A adhesin but not with control antibodies directed against Pil1. Results are means ± SD from 5 independent experiments performed in duplicate. Asterisks represent statistical differences relative to WT strain UCN34 with * p< 0.05; ** p<0.01; *** p<0.001 using two-way ANOVA with Bonferroni’s post-test in GraphPad Prism version 5.

## DISCUSSION

*S. gallolyticus subsp. gallolyticus* (*Sgg*) belongs to the *Streptococcus bovis/ Streptococcus equinus* complex, a diverse group of streptococci that are commensals of the gut, opportunistic pathogens and used in dairy product fermentation. *Sgg* is described as a weak colonizer of the the gastrointestinal tract with a fecal carriage of about 2.5 to 15%. It is believed that under certain specific physiological conditions such as development of colon malignancies, *Sgg* is able to overgrow by benefiting from specific tumoral nutrients and outcompeting closely related microbiota gut commensals (7, 8). This increase in *Sgg* load and the changes in the gut intestinal barrier resulting from tumor development are suspected to favor *Sgg* translocation across the tight intestinal barrier, which in turn can lead to invasive infections such as septicemia and infective endocarditis (19, 20).

In this work, we investigated the ability of *S. gallolyticus subsp. gallolyticus* strain UCN34 to translocate across intestinal barriers using Caco-2 and T84, two widely used model cell lines derived from human colon adenocarcinoma. Our results are in perfect agreement with a previous report by Boleij et al. showing that *Sgg* UCN34 could efficiently translocate across polarized Caco-2 cells while the closely related non-pathogenic *S. gallolyticus subsp. macedonicus* (*Sgm*) was not able to do so (3). Since *Sgm* does not possess any pili, we hypothesized that that the Pil3 pilus could be involved in translocation. Pili have long been considered important players in bacterial attachment to host tissues and their role in translocation across intestinal epithelia was elucidated for GBS (17).

Here, we demonstrate that translocation of *Sgg* UCN34 across polarized intestinal cells is a Pil3-dependent process. Indeed, the Δ*pil3* mutant was unable to translocate across Caco-2 and T84 monolayers. Expression of Pil3 in WT UCN34 is known to be heterogeneous at the population level, with a majority of cells weakly piliated (90%) and a minority highly piliated (10%). Interestingly we found that a Pil3+ variant homogeneously expressing high levels of Pil3 pilus is unable to translocate intestinal barriers. This result suggested that heterogeneous expression of Pil3 plays a key role in the translocation process. We were able to mimic this heterogeneity *in vitro* by mixing live Δ*pil3* mutant cells with the PFA-killed Pil3+ variant. Both the Δ*pil3* and Pil3+ variants were unable to translocate the intestinal barrier alone. Strikingly, when combined in a proportion similar to that found in the UCN34 WT population, about 9 Δ*pil3* for 1 Pil3+, we found that the presence of Pil3+ variant cells allowed translocation of the Δ*pil3* mutant. Based on these results we propose the following model to explain crossing of epithelial junctions by *Sgg*. Highly piliated Pil3 bacteria in the UCN34 population interact with an unknown cell surface receptor, or a component of the tight junctions, likely through the Pil3A adhesin, activating signaling pathway(s) involved in regulation of epithelial cell junctions. This then leads to opening of cell junctions allowing loosely bound bacteria with low Pil3 levels to pass in between adjacent cells. This translocation occurs without major disruption of the epithelial junctions. As shown for Pil1, this heterogeneity of Pil3 pilus expression can also mitigate host immune responses allowing *Sgg* to more efficiently evade host intestinal immune defenses in the lamina propria and later in the blood. Future studies will be aimed at investigating the identity of the Pil3A receptor on polarized cell monolayers.

## MATERIAL AND METHODS

### Cell cultures and bacterial strains

Caco-2 and T84 cells were routinely grown in Dulbecco’s Modified Eagle Medium (DMEM GlutaMAX, pyruvate and 4.5 g/L D-glucose) supplemented with 10% heat-inactivated fetal bovine serum (FBS). To obtain polarized Caco-2 and T84 monolayers, cells were trypsinized and seeded on inverted 12 mm polycarbonate, 3 μm-pore, tissue culture inserts (Transwell permeable support, Corning) at a density of 10^6^ cells/cm^2^. After incubation for 6 h at 37°C, transwell inserts were placed back into the wells and supplemented with fresh media every two days for 14 days. Trans Epithelial Resistance (TER) was measured with an Ohmmeter (Millicell-ERS, Millipore) and paracellular permeability was measured using the nonionic macromolecular tracer FITC-Dextran 4000 Da (Sigma). The cell medium in both compartments was removed. The lower compartment (corresponding to the apical surface of the epithelium) was replaced with RPMI without phenol red (Invitrogen) supplemented with FITC-dextran 4000 (5 mg/ml) and the upper compartment (i.e. basolateral side) with RPMI without phenol red. After incubation for 1 h, the upper compartment was sampled and absorbance at 490 nm measured. S. *gallolyticus* strains were grown at 37°C in Todd-Hewitt (TH) broth in standing filled flasks.

### Adherence assays

Caco-2 and T84 cells were seeded at 3 × 10^5^ cells ml^−1^ in 24-well plates and incubated at 37°C in 5% CO2 until 100% confluence. Overnight cultures of *S*. *gallolyticus* strains were washed once in PBS and resuspended in DMEM medium prior to infecting cells at a MOI of 10 bacteria per cell. Bacteria added to confluent monolayers were centrifuged at 500 RPM ≈90*g* to synchronize infections. After 2, 4, 6 h of incubation at 37°C under 5% CO2 atmosphere, monolayers were washed 4 times to remove non-adherent bacteria, cells were then lysed in cold water and plated to count cell-associated bacteria. The percentage of adherence was calculated as follows: (CFU on plate count / CFU in inoculum) X100. Assays were performed in triplicate and were repeated in at least 3 independent experiments.

### Bacterial translocation assays

Overnight cultures of *S. gallolyticus* were washed in PBS and resuspended at 1 × 10^8^ CFU/ml in pre-warmed DMEM. Cell monolayers were washed with DMEM and inverted in 6-well plates. 50μL of *Sgg* inoculum was then added to the apical side of the cells and incubated for 1h at 37°C in 5% CO2 atmosphere. The transwell inserts were placed back into the corresponding wells and fresh culture media was added to both compartments. At each time point of infection, the medium from the upper compartment was recovered for CFU determination and replaced by fresh media to prevent bacterial planktonic growth. For the preparation of the artificial mixture, *Sgg* Pil3+ cells were washed in PBS and fixed with 4% PFA for 20 min. Following fixation, bacteria were washed 4 times in DMEM. We verified that no CFU could be recovered after this treatment. Live Δ*pil3* mutant cells were then added to the PFA killed Pil3+ variant, a ratio of 9:1. For the blocking experiments with antibodies directed against Pil3, a combination of antibodies directed against the C- and N-terminal domains of Pil3A were added to the fixed Pil3+ variant and preincubated for 30 min at room temperature. This mixture was then added to the live Δ*pil3* mutant for cell monolayer infection as indicated above.

### Confocal microscopy

At 6 h post-infection, cell monolayers were washed once with PBS and then fixed with PFA 4% for 10 min at room temperature. Monolayers were then washed three times with PBS and subsequently quenched with Glycine 0.1M. For confocal microscopy, cells were permeabilized in PBS-Triton-X100 0.2% for 10 min at 4°C. Cell monolayers were then incubated with anti-UCN34 polyclonal antibody at a 1:200 dilution (Covalab, France), to specifically label *S*. *gallolyticus*, followed by incubation with secondary DyLight-488 conjugated goat anti-rabbit antibody (1:200). In addition, the adherens junction protein E-cadherin was stained using an anti-E-cadherin HECD-1 monoclonal antibody (Invitrogen) at 1:100 with subsequent incubation with the secondary DyLight594 conjugated anti-mouse antibody (1:100). Finally, Hoecht 33342 (1:2000) was added to visualize cell nuclei and Alexa Fluor 647 phalloidin (1:50) to detect the actin cytoskeleton. Samples were mounted using ProLong Gold Antifade reagent and Z-stacks of 300 nm step size were acquired using a Leica TCS SP5 confocal microscope with a 63x oil objective. Immunofluorescence images were analyzed using the Fiji software.

## Acknowledgments

We sincerely thank Tarek Msadek for the critical reading of the manuscript. This work was supported by the French Government’s “Investissement d’Avenir” program Laboratoire d’Excellence “Integrative Biology of Emerging Infectious Diseases” Grant ANR-10-LABX-62-IBEID. Mariana Martins was funded by the PPU program and the ARC foundation.

## Supplementary Figures Legend

**Fig. S1. Monitoring the epithelial integrity of Caco-2 monolayer.** A- Permeability of the monolayer to 4 kDa- FITC fluorescent dextran added in the upper compartment at 5 mg/ml for 1h. B- Measurement of transepithelial resistance (TER) after various days of culture. C- Measurement of the TER at the beginning of infection (NI) and after 6h of infection with *Sgg* UCN34. Results are means ± SD from 3 independent experiments performed in duplicate.

**Fig. S2. Adhesion and translocation of *Sgg* across T84 epithelial monolayer**. Adherence of UCN34, Pil3+ and Δ*pil3* is presented as percentage of bacterial inoculum after 4h and 6h at 37°C at a multiplicity of infection of 10 bacteria per cell. B- UCN34, Pil3+ and Δ*pil3* translocation across T84 monolayer after 2 h, 4 h and 6 h of infection. C- Monitoring of the epithelial integrity of T84 monolayers by permeability to 4 kDa FITC dextran and measurement of transepithelial resistance (TER) after various days of culture. D- Measurement of the TER at the beginning of infection (NI) and after 6h of infection with *Sgg* UCN34. Results are means ± SD from 3 independent experiments performed in duplicate. Note that the transepithelial resistance of T84 cells is 10-fold higher than in Caco-2 cells reaching 3,000 ohms per square centimeter after 11 days of culture.

